# FLYWCH transcription factors act in a LIN-42/Period autoregulatory loop during gonad migration in *C. elegans*

**DOI:** 10.1101/2025.07.10.664215

**Authors:** Brian Kinney, Jason Menjivar-Hernandez, Fred Koitz, Dmitry Grinevich, Madonna Baselios, Christopher M. Hammell, Kacy Lynn Gordon

## Abstract

Development must be coordinated across body systems but must also accommodate cell-type-specific processes. We discovered that the gene regulatory circuit controlling developmental timing in the *Caenorhabditis elegans* larval skin exhibits both points of convergence and divergence with the regulatory program governing the migration of the leader cell in gonad development, the distal tip cell (DTC). As a point of convergence, the conserved regulator of developmental timing, LIN-42/Period, peaks synchronously across cell types both during the L3 stage, when the DTC makes a turn in its normal migratory path, and the L4 stage in which the DTC normally continues straight ahead. We report that *lin-42*, like its ortholog *period*, autorepresses its own transcription. *lin-42* is required cell-autonomously for proper pathfinding of the DTC; DTC-specific *lin-42* RNAi causes the DTC to turn in the mid-L4 instead of continuing straight ahead. We identified the FLYWCH transcription factor FLH-1 as able to directly bind the *lin-42a* promoter. Using live-cell imaging, we show that *flh-1; flh-2* double mutant DTCs have an aberrant turn in the mid-L4. These mutants derepress the L4 peak of *lin-42* expression in a stage- and DTC-specific manner, and this derepression is itself *lin-42-*dependent. During the aberrant mid-L4 turn in *flh-1*; *flh-2* mutants, the focal adhesion factor TLN-1 is repolarized in the direction of turning. These results reveal that bodywide developmental rhythms can be fine-tuned to integrate with specific organogenic processes.

**SUMMARY STATEMENT:** To identify new transcriptional regulators of distal tip cell migration, a screen for *C. elegans* transcription factors that bind the promoter of the Period homolog *lin-42*, a conserved regulator of developmental timing, uncovers a pair of factors that redundantly prevent misdirected migration of the distal tip cell–the cell that guides gonadal development. Through phenotypic analysis of genetic mutants, RNAi knockdown, and live imaging, we found that these transcription factors create a cell-specific autoregulatory loop that controls *lin-42* transcription and promotes directional migration by polarizing the focal adhesion protein TLN-1.

## INTRODUCTION

Determining how cells within a developing organism are patterned in three-dimensional space is a major focus of developmental biology research. How development is coordinated across cell types in the fourth dimension–time–is comparatively less well understood. Genetic studies of the nematode *Caenorhabditis elegans* have been pivotal for the understanding of developmental timing. Heterochronic mutants, which either repeat or skip larval developmental events, shed light on the genetic regulation of developmental timing and led to the discovery of microRNA-based gene regulation (Ambros, 2011; Ambros and Horvitz, 1984). The heterochronic microRNA gene *lin-4* regulates the molting cycle—encompassing the divisions and fusions of the larval skin cells and the shedding and synthesis of new cuticles—across the four larval stages (Ambros, 2011; Ambros and Horvitz, 1984; Chalfie, 1981). *lin-4* mutants repeat larval cell division patterns (Chalfie, 1981) and fail to undergo certain morphogenetic events, such as vulva formation (Chalfie, 1981; Euling and Ambros, 1996). Recent work demonstrated that *lin-4* transcription is pulsatile at the midpoint of each larval stage (Kinney et al., 2023). Central to the *lin-4* regulatory network is the Period homolog LIN-42, which represses *lin-4* expression at each molt to help temporally refine *lin-4* expression (Perales et al., 2014). LIN-42 acts through *lin-4* co-activators NHR-23/ROR and NHR-85/Rev-Erb to repress *lin-4* (Kinney et al., 2023). The *lin-42* gene also regulates gonad development (Tennessen et al., 2006). We asked whether the newly described transcriptional regulatory circuit that controls molting also regulates gonad development, and if not, what other factors work with *lin-42* as a point of convergence.

Gonad development proceeds in a linear sequence of events. The germline stem cell niche and leader cell of gonad elongation is the distal tip cell, or DTC (Gordon, 2020; Hubbard and Greenstein, 2000; Kimble and White, 1981), and its migration is powered in part by pushing forces from the proliferating germ cells (Agarwal et al., 2022), in addition to a germ cell-independent force yet to be identified (So et al., 2024). Gonad elongation begins with each DTC leading each of the two arms of the hermaphrodite gonad away from the ventral midbody in the L2 larval stage. The DTC makes two 90-degree turns in the L3 larval “turning” stage and elongates back to the dorsal midbody during the L4 larval “pathfinding” stage to create the two U-shaped arms of the adult gonad (Hedgecock et al., 1987; Hubbard and Greenstein, 2000; Kiyoji Nishiwaki, 1999). We probed potential overlap and divergence between the mechanisms regulating developmental timing in the larval DTC vs. the larval skin.

Knockdown of *lin-42* results in precocious DTC turning (Tennessen et al., 2006). While antibody staining detected LIN-42 in the DTC during the L2 and L3 stages, it was undetectable in that cell during the L4 stage (Tennessen et al., 2006). This suggests a L4-specific mechanism to repress LIN-42 in the DTC during the L4 pathfinding stage.

We hypothesized that the larval developmental timing network has points of overlap that keep tissues developing in sync and points of divergence that allow for differences in developmental trajectory across cell types. We aimed to investigate how timing factors operate in the DTC, and which other factors may regulate timing in this cell that leads gonad morphogenesis.

Here, we report that the factors that work with *lin-42 to* regulate timing in the seam cells, *lin-4,* NHR-23 and NHR-85, are not necessary for DTC migration. DTC-specific RNAi of *lin-42* causes a DTC pathfinding defect we call a “ventral return”. Furthermore, *lin-42* is autorepressive in that and other cell types. We identified FLH-1 as a potential transcriptional regulator of *lin-42* by conducting a yeast one-hybrid screen using a *lin-42* promoter as bait against 937 predicted *C. elegans* transcription factors (Reece-Hoyes et al., 2011) and found that it represses *lin-42* expression redundantly with a partner *flh-2*. In a *flh-1; flh-2* double hypomorphic mutant, the DTC also makes a ventral return during the L4 pathfinding stage. The ventral return is associated with mislocalization of GFP::TLN-1, a protein that crosslinks integrin INA-1 and PAT-3 to the actin cytoskeleton and is required for proper DTC migration (Agarwal et al., 2022; Wong et al., 2014). Genetic evidence indicates that FLH-1/2 are transcriptional co-repressors that function alongside (and repress the transcription of) *lin-42* in the DTC. Our results show that *lin-42* expression across cell types peaks and falls in synchrony, yet specific developmental events depend on cell-and stage-specific refinements of *lin-42* regulation. In the DTC, *flh-1* and *flh-2* negatively regulate *lin-42* expression to ensure proper migration of the DTC during the final phase of gonad elongation.

## RESULTS

### The developmental timing system in the larval skin intersects with DTC regulation at *lin-42,* which is required for DTC pathfinding in the L4 stage

The *C. elegans* ortholog of Period, *lin-42*, regulates both the timing of DTC turning during the L3 stage (Tennessen et al., 2006; Cecchetelli and Cram, 2017) and the molting cycle of the larval skin (Kinney et al., 2023). We investigated whether the same gene regulatory network governs developmental timing across larval stages in both of these developmental programs. We imaged endogenously tagged LIN-42::YFP along with its known partners in the larval skin and vulva, NHR-23::mScarlet and NHR-85::GFP (Kinney et al., 2023). We examined seam cells, hyp7 hypodermal cells, vulval cells, and the DTC across the L3 and L4 substages until young adulthood using P6.p cell number and vulval morphology to identify each substage (Fig. 1A, B) (Mok et al., 2015). While NHR-23::mScarlet and NHR-85::GFP exhibited synchronized dynamic expression during the L3 and L4 substages, peaking at L4.3 and L4.9 respectively across the seam cells, hyp7 cells, and the vulval cells, neither was detected above background in the DTC at any stage (Fig. 1C-F). Neither *nhr-23* RNAi nor *nhr-85(ok2051)* loss of function had a notable DTC migration defect rate (Fig. S1). In contrast, LIN-42::YFP peaks in the DTC synchronously with other examined cell types in the middle of L3 and L4 (Fig. 1B-F), consistent with its previously published expression dynamics (Jeon et al., 1999; Monsalve et al., 2011). Previous immunohistochemistry did not detect LIN-42 in the L4 DTC (Tennessen et al., 2006). LIN-42 expression levels in the DTC are notably lower than in other cell types (Fig. 1C-F), so this discrepancy is likely a difference in sensitivity.

**Figure 1.**
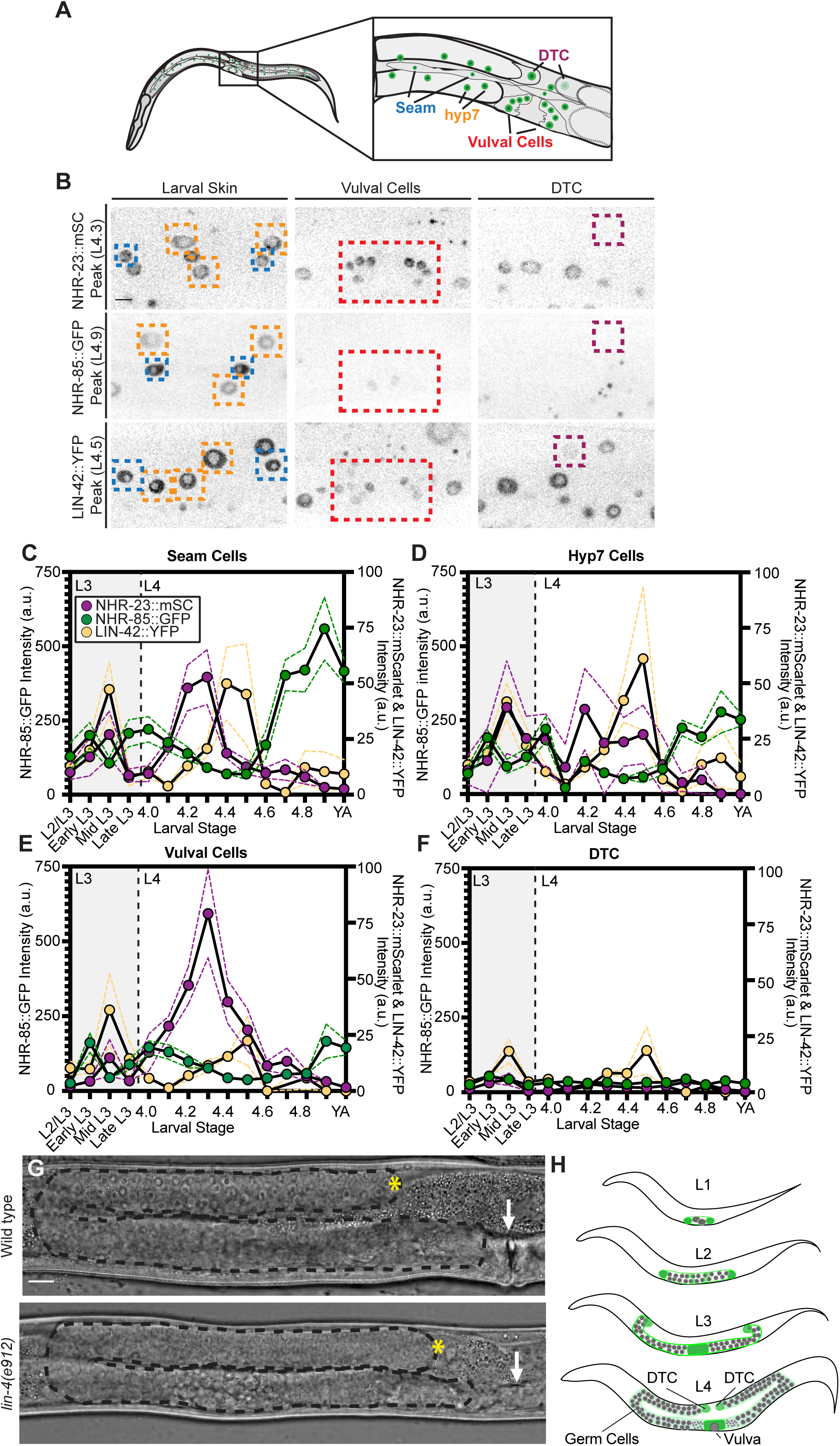
Dynamic LIN-42 expression is synchronized across the larval skin and DTC during larval development. (A) Schematic showing the locations of the different cell types analyzed. (B) Fluorescence images of NHR-23::mScarlet, NHR-85::GFP, and LIN-42::YFP at peak expression across the cell types measured (marked with a colored dotted box), which include hypodermal seam cells (blue) and hyp7 (orange), vulval cells (red), and DTC (purple). (C-F) Quantification of NHR-85::GFP (green), NHR-23::mScarlet (purple), and LIN-42::YFP (both A and B isoforms tagged, yellow) expression levels in (C) hypodermal seam cells, (D) hyp7 cells, (E) vulval cells, and (F) DTC across the L3 and L4 substages. Each data point represents the fluorescence intensity from an individual animal measured as the mean intensity of three cells per cell type (seam, hyp7, vul), or intensity of a single DTC per animal (n=8-12 per substage). Dotted lines show SEM. (G) Comparison of *lin-4(e912)* mutant to wild type animal 36 hours after L1 synchronization. The site of the vulva (or the absence of one) is marked by a white arrow, the gonad is outlined with a dotted black line, and the DTC is marked by a yellow asterisk (*). Wild type n=34, *lin-4(e912)* n=30. Scale bar: 10 μm. (H) Schematic illustrating *C. elegans* gonad development through the larval stages. The DTCs and proximal gonad are shown in green, and germ cells are shown in gray.

The peak of LIN-42 expression in the DTC distinguishes it from NHR-85 and NHR-23 (Kinney et al., 2023), which are absent from that cell across the stages examined (Fig. 1C-F). Furthermore, the known target of these regulatory factors in the larval skin, the microRNA gene *lin-4,* is dispensable for gonad morphogenesis (Fig. 1G) (Ambros, 1989; Ambros and Horvitz, 1984; Bracht et al., 2010). *lin-4(e912)* mutants are fertile with grossly normal gonad morphology (though they lack vulva formation) (Chalfie, 1981; Euling and Ambros, 1996). The DTC therefore employs a different genetic timing mechanism than that which governs the larval skin (Fig. 1H), but these cell types share dynamic *lin-42* expression.

Maternally injected whole-body RNAi to *lin-42* is known to cause precocious DTC turning during the L2 stage in a minority of animals (Tennessen et al., 2006). However, whether or how *lin-42* regulates the DTC during the L4 pathfinding stage is not known. To investigate a potential role for *lin-42* in the L4 DTC, we constructed an RNAi clone targeting both major *lin-42* isoform mRNAs (Methods, Fig. S2), which causes known *lin-42* loss of function phenotypes like precocious alae (Banerjee et al., 2005) and precocious DTC turning (Tennessen et al., 2006), and quantitative knockdown of LIN-42::YFP of 74% in hyp7 cells (Fig. S2A-D). To assess the role of *lin-42* in the DTC, we used a DTC-specific RNAi strain with a DTC mNeonGreen membrane marker (Linden et al., 2017). First, *lin-42* RNAi treatment on the DTC-specific strain displayed precocious turns in 2/31 animals (Fig. S2C), demonstrating that this previously observed *lin-42* loss-of-function phenotype is caused cell-autonomously by loss of *lin-42* in the DTC. Mistiming of the DTC turn in a minority of animals does not affect our ability to assess later L4 DTC migration defects, as both early-turning and normal-turning gonads have migrated to the dorsal body wall by mid-L4. After DTC-specific (Fig. 2A, B) *lin-42* RNAi, we observed a phenotype we dub “ventral return” in 24% (n=12/50) of L4.5-young adult animals. In these worms, the DTC dives sharply off the dorsal body wall and completes its migration along the ventral body (Fig. 2A). We conclude that *lin-42* is required in the DTC for proper pathfinding at the end of migration.

**Figure 2.**
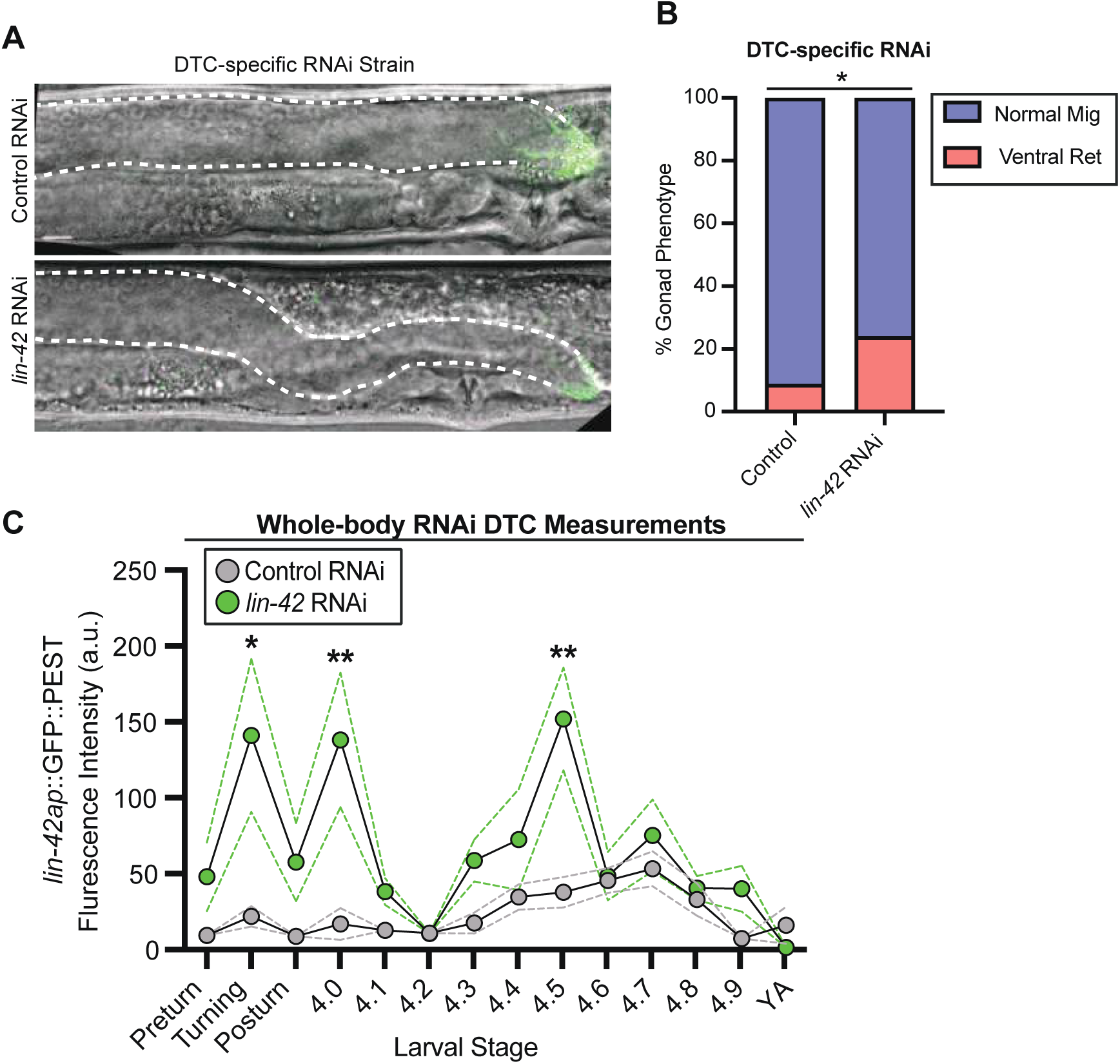
*lin-42* acts in an autoregulatory loop that regulates the DTC during the L4 pathfinding stage. (A) Representative images of animals treated with *lin-42* RNAi in a DTC-specific RNAi strain. The DTC is shown in green, and the dorsal gonad arm is outlined with a white dotted line. Scale bar: 10 μm. (B) Quantification of the ventral return phenotype comparing control (n=57) and *lin-42* RNAi treated (n=50) animals using the DTC-specific RNAi strain. Statistical analysis was performed with a two-tailed Fisher’s exact test (*p <0.05). (C) Expression intensity of *lin-42ap*::*SV40-NLS::GFP::PEST* in wild type and *lin-42* whole body RNAi treated animals in the DTC during each L3 turning stage and each L4 substage (n=8-12 per substage). Dotted lines show SEM. Two-way ANOVA with Tukey’s multiple comparisons test (F (27, 217) = 14.91) (**p <0.01)(*p <0.05).

### Dynamic expression of LIN-42 is transcriptionally autorepressed via the *lin-42a* promoter

We next asked how *lin-42* is transcriptionally regulated in the DTC. We generated two transcriptional reporters for *lin-42* using the putative promoters of *lin-42a* and *lin-42b/c* (hereafter referred to as *lin-42b*) (Edelman et al., 2016) to drive the expression of a nuclear-localized fluorescent protein with a PEST tag that promotes protein turnover (Fig. S3A-C). We observe dynamic expression of both *lin-42ap*::*SV40-NLS::GFP::PEST* (hereafter abbreviated *lin-42a::GFP*) and *lin-42bp::SV40-NLS::mCherry::PEST* (hereafter abbreviated *lin-42b::mCherry*) in the seam, hyp7, vulval cells, and the DTC (Fig. S3D). Transcription of *lin-42ap::GFP* rises just before the peak of LIN-42::YFP protein signal in the seam, hyp7, vulval cells, and DTC (shown in Fig. 1D-F). Transcription of *lin-42bp*::*mCherry* peaks after the timing we observe for the LIN-42::YFP peak (Fig. S3D). Transcripts of *lin-42a* and *lin-42b* both oscillate synchronously in L3–L4 staged bulk RNA-seq data (Fig. S4A, B) (Hendriks et al., 2014; Kim et al., 2013).

We focused on the *lin-42a* promoter because its transcription dynamics more faithfully anticipate the LIN-42::YFP protein signal dynamics observed synchronously across tissues (Fig. S3D). Furthermore, previous research indicates that the *lin-42(n1089)* deletion allele, which affects exons only in the *lin-42b/c* isoforms, has no DTC turning defects, whereas RNAi against the shared exons of *lin-42a/b* isoforms causes premature turning (Tennessen et al., 2006).

*lin-42* is the *C. elegans* ortholog of *Drosophila melanogaster* gene *period* (Tennessen et al 2006), the oscillation of which sets the 24-hour circadian cycle of the fly (Zehring et al., 1984; Bargiello et al., 1984). Clock gene oscillation is conserved in animals, including humans (Takahashi, 2017). It was discovered that *period* oscillation arises from negative transcriptional autoregulation of the *period* gene by the PER protein (Hardin et al., 1990; Young, 1996). To test for *lin-42* autoregulation, we treated the *lin-42* transcriptional reporter strain with *lin-42* RNAi (Fig. 2C). Whole-body RNAi knockdown of *lin-42* on the reporter strain resulted in overexpression of *lin-42ap::GFP* relative to the control in the DTC (Fig. 2C), as well as in hyp7 (Fig. S5; the same is observed for *lin-42b::mCherry::PEST* in hyp7 (Fig. S5). This derepression is observed during the mid-L3 stage when turning occurs, around the L3/L4 molt, and in the mid-L4 stage, when the ventral return is seen. *C. elegans lin-42* negatively autoregulates its own transcription, like the *period* gene of *Drosophila.* LIN-42 is not thought to bind DNA directly but instead works through binding partners (Lamberti et al., 2024; Kinney et al., 2023). Since dynamic *lin-42a* expression appears to be a point of convergence between the developmental timing network in the DTC and the rest of the body, we set out to investigate transcription factors that can bind the upstream regulatory region of *lin-42a*.

### FLH-1 can interact with the *lin-42a* promoter

To identify potential transcription factor candidates that may regulate *lin-42a* expression in the DTC, we performed a yeast one-hybrid (Y1H) screen. We constructed a yeast strain bearing a 1,515-bp putative *cis-*regulatory fragment from upstream of the *lin-42a* start site as bait (Fig. 3A) and mated this strain with every strain of a yeast library containing 937 predicted *C. elegans* transcription factors (Deplancke et al., 2006; Reece-Hoyes et al., 2011). We screened for interactions to identify candidate genes, which we then assessed for DTC phenotypes after loss-of-function in worms.

**Figure 3.**
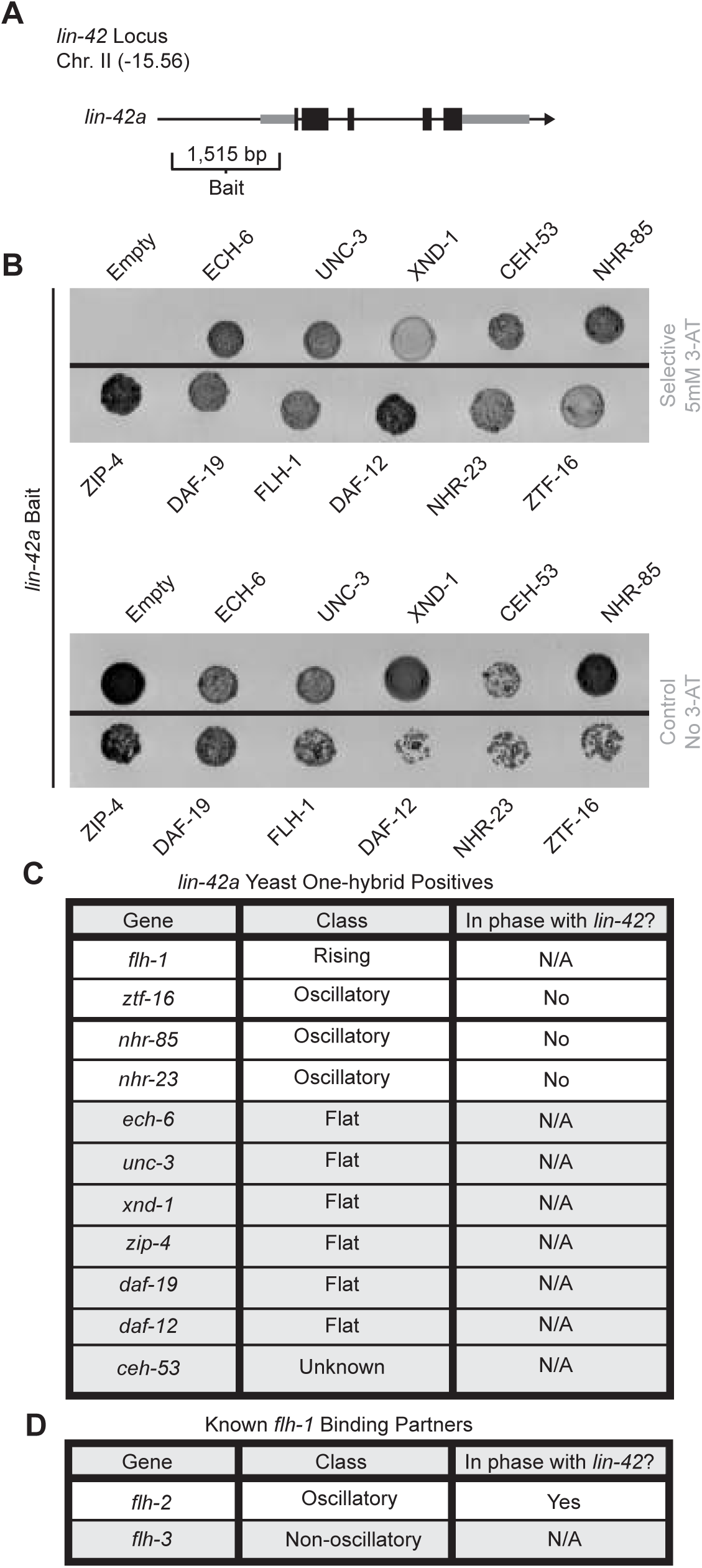
A yeast one-hybrid screen identifies potential transcriptional regulators of *lin-42*, one of which has a partner that oscillates in phase with *lin-42*. (A) Diagram of the *lin-42* locus and the region used as bait in the yeast one-hybrid screen. (B) Growth of strains positive for binding transcription factors at the *lin-42a* bait region on 5 mM 3-AT (top) and control plates without 3-AT (bottom). (C) List of *lin-42a* bait-binding transcription factor genes, including expression classification and phase (from Hendriks et al., 2014 and Meeuse et al., 2020). (D) List of genes encoding FLYWCH factors known to bind to FLH-1 including expression classification and phase (from Hendriks et al., 2014 and Meeuse et al., 2020).

Our Y1H screen identified 11 transcription factors that bind the *lin-42a* promoter (Fig. 3B). Since transcripts of *lin-42* itself oscillate (Fig. S4A-B reanalysis of Hendriks et al., 2014), we hypothesized that the dynamic expression of *lin-42a* might result from coregulation by a dynamically expressed transcription factor. Cross-referencing the list of 11 candidates with results from studies of oscillatory transcripts in *C. elegans* (Hendriks et al., 2014; Meeuse et al., 2020) reveals that most potential *lin-42a* promoter-binding transcription factors exhibit flat transcriptional profiles (Fig. 3C). Of those with dynamic expression, we have already ruled out NHR-23 and NHR-85 as regulators of DTC migration (Fig. 1F). The potential for direct binding of NHR-23 and NHR-85 to the *lin-42a* promoter was not previously known. Next, we find *ztf-16,* which is known to regulate DTC specification (Large and Mathies, 2010). Like *lin-42*, *ztf-16* prevents precocious expression of the adult-specific collagen gene *col-19* (Hansen et al., 2022; Abrahante et al., 1998). Finally, the Y1H candidate *flh-1* has a transcriptional profile that rises across larval stages (Fig. S4C), and the gene encoding its known partner, *flh-2* (Ow et al., 2008), oscillates in phase with *lin-42* (Fig. 3D) (Hendriks et al., 2014; Meeuse et al., 2020). A gene encoding another known binding partner of *flh-1*, *flh-3* (Ow et al., 2008), is also non-oscillatory (Fig. 3D) (Meeuse et al., 2020).

*flh-1* encodes a FLYWCH transcription factor that is known to cooperatively repress embryonic expression of the heterochronic microRNA *lin-4* with another FLYWCH transcription factor, FLH-2; mutants of these genes do not have postembryonic heterochronic defects in seam cell divisions or *col-19* expression (Ow et al., 2008). Yeast two-hybrid assays have detected an interaction between FLH-1 and FLH-2 (Reece-Hoyes et al., 2013), and transcriptomic evidence indicates that both *flh-1* and *flh-2* are expressed in adult DTCs (Ghaddar et al., 2023; Kaletsky et al., 2018). We do not observe evidence of FLH-2 directly binding the *lin-42a* promoter in the Y1H assay (Fig. 3B). These FLYWCH transcription factors have no known regulatory relationships with *lin-42*, nor any known role in DTC migration, which we set out to investigate.

### *flh-1* and *flh-2* redundantly regulate DTC migration and *lin-42* expression in the L4 stage

To determine whether these FLYWCH transcription factors are functionally implicated in DTC migration, we investigated FLYWCH mutant young adult gonads for evidence of DTC migration defects. None of the *flh-1, flh-2,* or *flh-3* single mutants had a migration defect significantly different from that of the wild-type strain, although we did observe animals with the ventral return phenotype (Fig. S1). We next tested whether *flh-1* and *flh-2* might redundantly co-regulate DTC migration and enhance the low-penetrance defect we observed in the single mutants (Fig. S1). Since *flh-1(bc374)*; *flh-2(bc375)* double mutants die as young larvae (Ow et al., 2008), we tested a double mutant with the hypomorphic *flh-1(tm2118)* (a 707-bp deletion) and *flh-2(tm2126)* (a 348-bp deletion) alleles (Ow et al., 2008). From here on, we refer to this as the *flh-1/2* mutant. Single mutants of *flh-1(tm2118)* and *flh-2(tm2126)* only had 6.45% and 6.25% penetrance of the ventral return phenotype, respectively compared to the 6% occurrence in wild-type animals (Fig. 4A, B, Fig. S1D). However, *flh-1/2* double mutants displayed a greater penetrance (28.85%) (n = 15/52) of the ventral return phenotype (Fig. 4A, B). We performed RNAi knockdown for *flh-3* on *flh-1/2* double mutant worms but saw no enhancement (21.875% penetrance of the ventral return phenotype, Fig. S1D). We conclude that *flh-1* and *flh-2* redundantly regulate DTC migration by preventing ventral return during the L4 stage.

**Figure 4.**
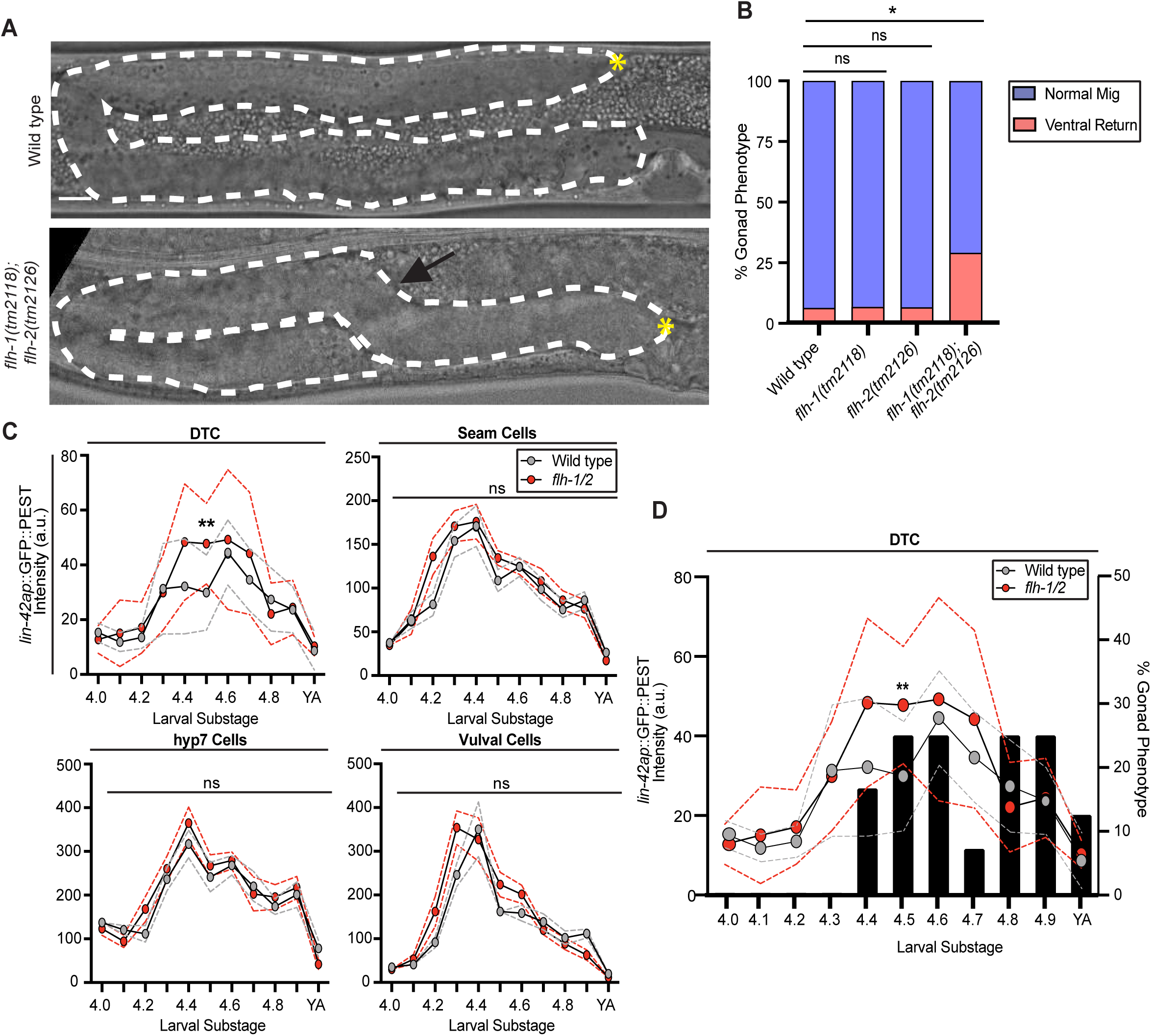
*flh-1* and *flh-2* hypomorphic double mutants display defects in migration and *lin-42* transcription in the L4 DTC. (A) Wild type (top) and *flh-1(tm2118)*; *flh-2(tm2126) (flh-1/2)* mutants (bottom) at the late L4 stage. The gonad is outlined by a dashed whited line and the DTC is marked by a yellow asterisk (*). *flh-1/2* double mutant animals display a “ventral return” (black arrow). Scale bar: 10 μm. (B) Percentage of samples with each phenotype for genotypes shown at bottom. (Wild type n=50, *flh-1(tm2118)* n=31, *flh-2(tm2126)* n=32, *flh-1/2* double mutant n=52,. Statistical analysis was performed using a pairwise proportion test, with p-values adjusted for multiple comparisons via the Benjamini-Hochberg procedure (*p <0.05). (C) Expression intensity of *lin-42ap::SV40-NLS::GFP::PEST* in wild type (gray)) and *flh-1/2* double mutant (red) animals in the DTC, seam cells, hyp7 cells, and vulval cells in each L4 substage. Each data point represents the fluorescence intensity from an individual animal measured as the mean intensity of three cells per cell type (seam, hyp7, vul), or intensity of a single DTC per animal (n=8-12 per substage). Dashed lines represent SEM. Statistics were performed using a Two-way ANOVA with Tukey’s multiple comparisons test (F (21, 184) = 11.31) (**p <0.01). (D) Bar graph displaying the percentage of samples with the ventral turn phenotype in *flh-1/2* double mutant animals at each L4 substage using the images quantified and overlaid with *lin-42ap::SV40-NLS::GFP::PEST* expression in the DTC from (C) (n=8-14 per substage).

To test for regulation of *lin-42* transcription by *flh-1/2 in vivo,* we used our *lin-42* transcriptional reporters to compare the expression of the *lin-42a* promoter in the wild type and *flh-1/2* double mutant in the seam cells, hyp7, vulval cells, and the DTC. Comparing *lin-42a* transcriptional expression in otherwise wild-type worms to reporter expression in a *flh-1/2* double mutant background showed significant derepression of the reporter at the peak of expression only in the DTC, during the L4.5 substage (Fig. 4C). The earliest appearance of the ventral return phenotype coincided with the divergence of *lin-42a* transcription between *flh-1/2* mutants and controls (Fig. 4D). We hypothesize that FLH-1 and FLH-2 work redundantly to attenuate peak expression of *lin-42a* selectively in the DTC at the mid-L4 stage. This proposed regulatory interaction concords with the observation that *lin-42* and *flh-2* transcription oscillates in the same phase (Fig. 3D) (Hendriks et al., 2014; Meeuse et al., 2020). Expression driven by *lin-42b* was also altered in the *flh-1/2* double mutant background (Fig. S6), though neither FLH-1 nor FLH-2 bind this region in yeast one-hybrid (Fig. S7). The genetic evidence demonstrates that *flh-1*/*2* mutants misregulate *lin-42* expression levels and the pathfinding stage of migration in the DTC.

### The *lin-42* autoregulatory loop includes *flh-1*/*2*

To test whether the FLYWCH factors help mediate the autoregulation of the *lin-42* promoter that we observed in Fig. 2C, we exposed wild-type control animals and *flh-1/2* mutant animals to empty vector L4440 and *lin-42* RNAi (Fig. 5A). After whole-body *lin-42* RNAi of wild-type animals, we measured the ventral return phenotype in n=17/71, or 23.94% of animals (Fig. 5A). *flh-1/2* mutants on control RNAi have a similar frequency of the ventral return phenotype (n=15/57, 26.32%). Indeed, these two defect rates are not significantly different from one another. However, they both differ from wild-type worms on control L4440 RNAi vector (n=5/69) (Fig. 5A). Importantly*, flh-1/2* double mutants on *lin-42* RNAi had neither enhancement nor suppression of the ventral return phenotype (Fig. 5A), suggesting *flh-1/2* and *lin-42* act in the same pathway and in the same direction.

**Figure 5.**
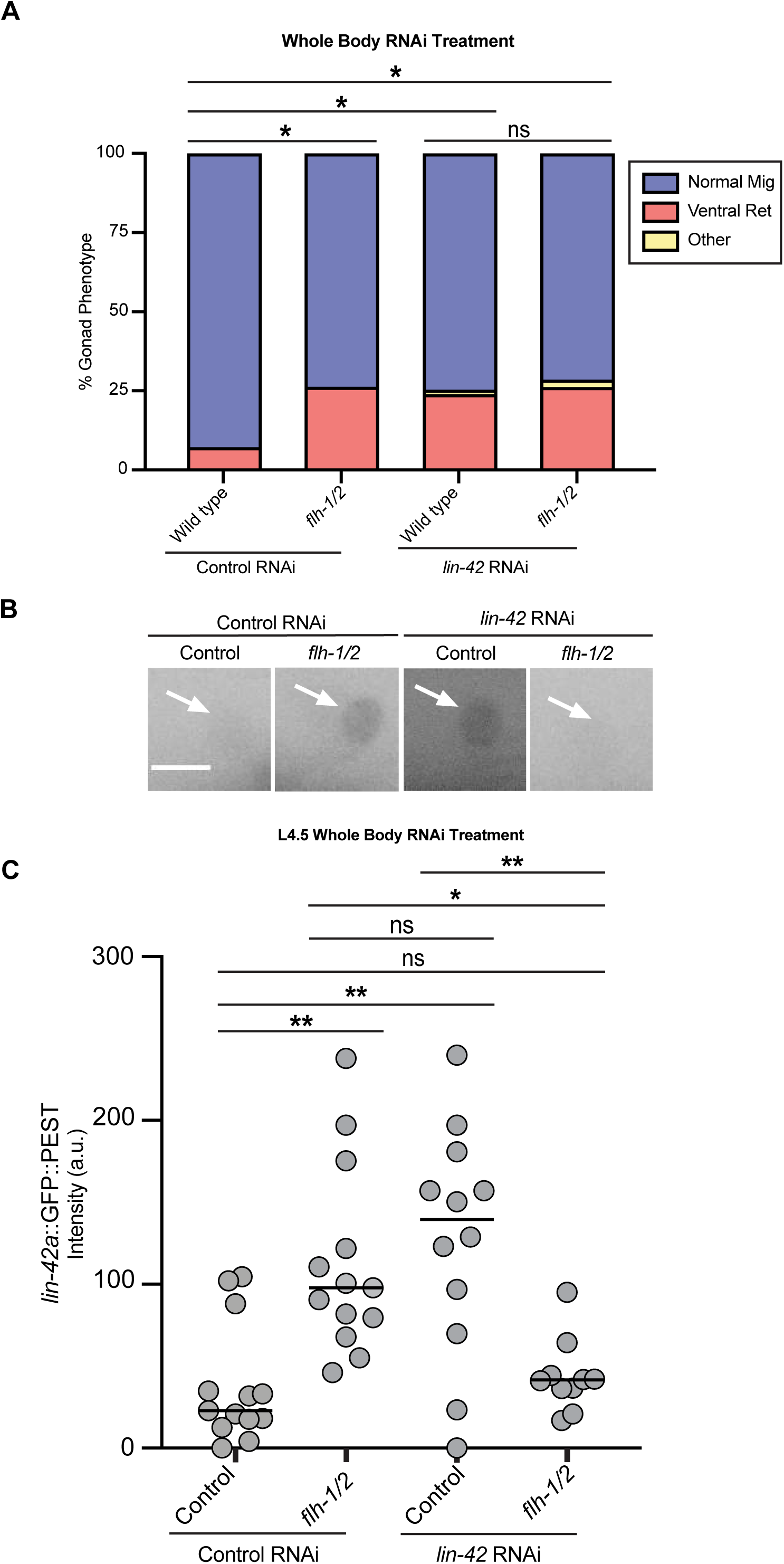
*flh-1/2* regulate *lin-42* as part of an autoregulatory loop. (A) Percentage of samples with each phenotype for conditions shown at bottom. (Control RNAi of wild type n=69, control RNAi of *flh-1/2* n=57, *lin-42* RNAi of wild type n=71, *lin-42* RNAi of *flh-1/2* n=42). Each animal was classified as Normal Migration (blue), Ventral Return (red), or Other (yellow). Statistical analysis was performed using a pairwise proportion test for number of animals with ventral return phenotype, with p-values adjusted for multiple comparisons via the Benjamini-Hochberg procedure (*p <0.05). (B) Inverted representative images of *lin-42ap::SV40-NLS::GFP::PEST* in L4.5 DTCs comparing control RNAi and *lin-42* RNAi in wild type and *flh-1/2* double mutant backgrounds. Scale bar: 5 μm. (C) Expression intensity of *lin-42ap*::*SV40-NLS::GFP::PEST* in L4.5 DTCs comparing wild-type animals on control RNAi (gray, n=23), wild-type animals on *lin-42* RNAi (blue, n=16), *flh-1/2* double mutants on control RNAi (red, n=18), and *flh-1/2* double mutants on *lin-42* RNAi (yellow, n=20). Wild type on control RNAi vs *flh-1/2* on control RNAi replicate of the results of (Figure 4C) L4.5. Some L4.5 animals for both control RNAi and lin-42 RNAi treated animals were the same samples used in (Fig. 2C). Two-way ANOVA with Tukey’s multiple comparisons test (F (3, 51) = 5.346) (**p <0.01) (*p <0.05).

We next sought evidence about the molecular basis of the interaction between *flh-1/2* and *lin-42*. Both FLYWCH factors (Ow et al., 2008) and LIN-42 (McCulloch and Rougvie, 2014; Perales et al., 2014; Van Wynsberghe et al., 2014) are transcriptional repressors, and our yeast one-hybrid demonstrates that FLH-1 can bind the *lin-42a* promoter. Revisiting the L4.5 stage at which both *flh-1/2* double mutants (Fig. 4) and *lin-42* RNAi (Fig. 2) cause derepression of the *lin-42a* promoter, we measured promoter expression in the otherwise wild-type *lin-42a::GFP* reporter strain and the *flh-1/2* mutants carrying the reporter, on both control and *lin-42* RNAi. As expected, loss of function of the FLYWCH factors or LIN-42 caused derepression of *lin-42a*. Surprisingly, placing the *flh-1/2* mutants on *lin-42* RNAi did not cause derepression, but instead reverted *lin-42a* expression to its baseline level in the DTC (Fig. 5B, C), but not in hyp7 (Fig. S5C). Such reciprocal sign epistasis has been observed when multiple mutations affect genes organized in a signalling cascade, specifically when repressors are present (Nghe et al., 2018) and in cases where each single mutation enhances gene expression (Baier et al., 2023), as reviewed in Zhang et al. (2024).One interpretation of these results is that *lin-42* is not autorepressive in the *flh-1/2* double mutant background.

Furthermore, because RNAi knockdown of *lin-42* does not rescue DTC migration in the *flh-1/2* mutants, we conclude that *flh-1/2* also must have other targets that regulate the DTC ventral return in a *lin-42*-independent manner. Note that while expression of the exogenous *lin-42a* promoter transgene is reverted to baseline after *lin-42* RNAi treatment of the *flh-1/2* double mutant, *lin-42* gene product levels cannot be rescued because the experimental treatment of *lin-42* RNAi causes gene knockdown.

### *flh-1/2* repression of *lin-42* in the DTC is stage-specific and not observed in L3

We were intrigued to see *lin-42a* derepression after *lin-42* RNAi during the normal L3 turn (Fig. 2C, also note derepression at L4.0), and we asked if L3 *lin-42* autorepression was also co-regulated by FLH-1/2. During the L3 larval stage, the peak of *lin-42ap::GFP* expression coincides with the second stage of DTC migration, during which the DTC moves from the ventral body wall up to the dorsal body wall through two 90-degree turns (Fig. 6A). Expression of *lin-42a::GFP* is barely detectable before or after the turn (Fig. 6A-B). The pattern we observe in *flh-1/2* double mutants is qualitatively the same (Fig. 6A-B), and these mutants always execute the L3 turns normally (n=71/71 mid-L4-young adult *flh-1/2* mutants had normal U-shaped gonad bends). However, *lin-42a* reporter expression was weaker in the double mutants during the L3 turn (Fig. 6A-B; the same was observed for *lin-42b::mCherry*, Fig. S6). These results suggest that FLH-1 and/or FLH-2, potentially indirectly and/or with different binding partner(s), are *lin-42* activators rather than repressors in the L3 stage. *flh-1* expression in the whole body is low in the L3 stage (Fig. S4C).

**Figure 6.**
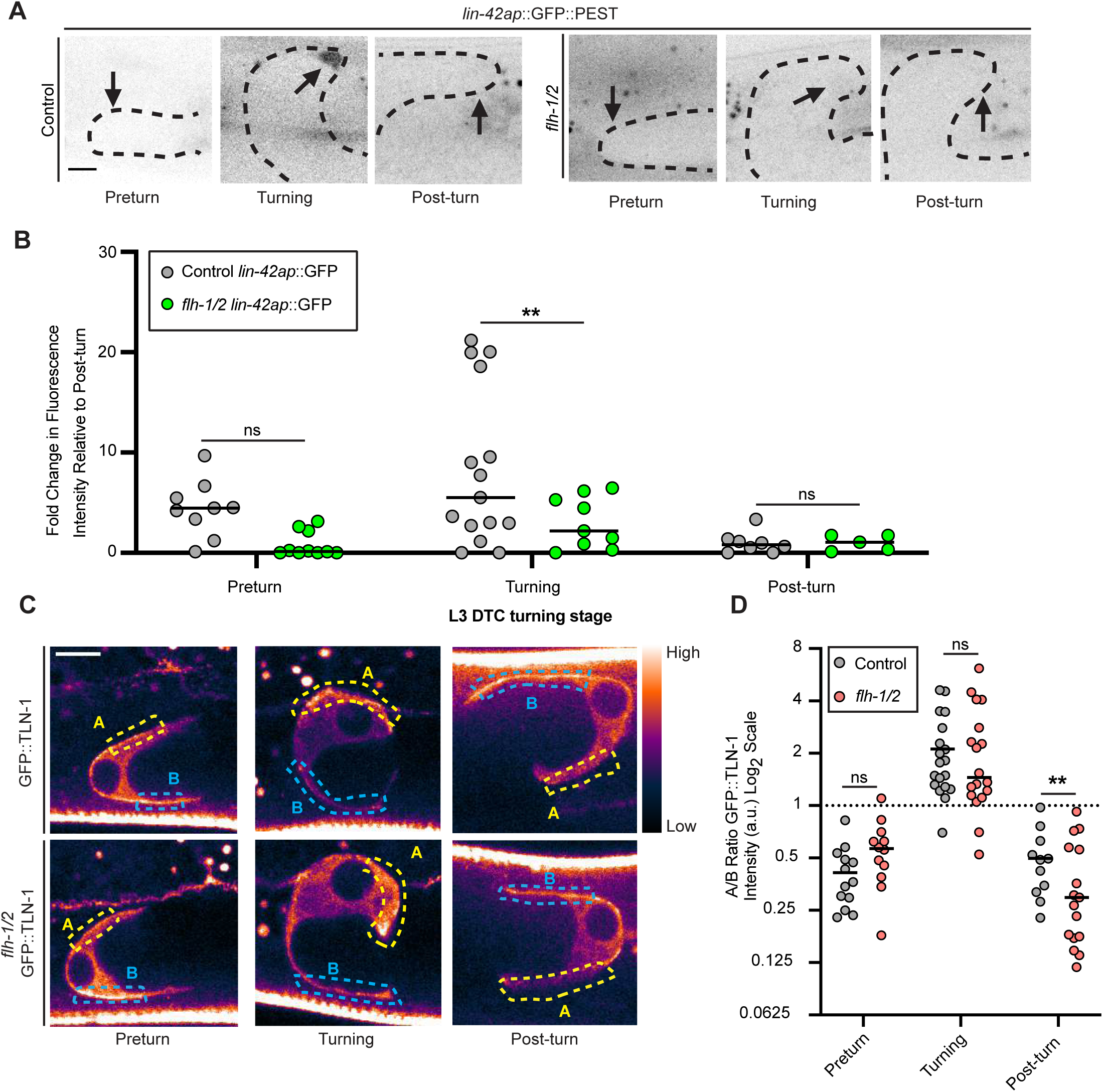
*flh-1/2* mutant DTCs have normal GFP::TLN-1 polarization during turning but less *lin-42a* expression. (A) Inverted representative images from three staged specimens each of *lin-42ap::SV40-NLS::GFP::PEST* in otherwise wild-type controls (left) and *flh-1/2* mutants (right) in the L3 DTC during pre-turning, turning, and post-turning stages. The gonad arm is outlined by a black dotted line and DTC nucleus marked with a black arrow. Scale bar 5 μm. (B) Quantification of fluorescence intensity in *lin-42ap::SV40-NLS::GFP::PEST* in otherwise wild-type controls (gray), *flh-1/2* (green), during each L3 DTC turning stage. Values are normalized to the mean post-turn expression in the dataset. For wild type: preturn n=9, turning n=15, and post-turn n=8. For *flh-1/2* mutants: preturn n=10, turning n=9, post-turn n=5. Two-way ANOVA with Tukey’s multiple comparisons test (F (5, 36) = 9.446) (**p <0.01). (C) Representative images of GFP::TLN-1 localization comparing control and *flh-1/2* double mutant DTCs in preturn, turning, and post-turn stages. The “gem” lookup table is used to show relative fluorescence intensity. Blue dotted box indicates the area of measurement of pre-turn ventral side of the DTC labeled “B”; the yellow dotted box shows the opposite cell surface measured labeled “A”. Scale bar: 5 μm. (D) Quantification of enrichment as ratio of A/B of intensity in GFP::TLN-1 comparing *flh-1/2* (red) to controls (gray) L3 DTCs at preturn, turning, and post-turn stages. The dotted line y = 1 (equivalent intensity) is provided for reference. For wild type: preturn n=13, turning n=19, post turn n=11. For *flh-1/2* mutants: preturn n=12, turning n=18, post-turn n=19. Two-way ANOVA with Tukey’s multiple comparisons test (F (5, 68) = 19.31) (**p <0.001).

Recent work has established that the DTC interacts with the extracellular matrix, in part through integrin-based adhesions, to regulate the turning phase of migration (Agarwal et al., 2022; Singh et al., 2024). TLN-1/talin is essential for proper DTC migration (Wong et al., 2014) and polarizes during DTC turning (Agarwal et al., 2022). This protein links integrins to the cytoskeleton (Waterston and Barnstead, 1996) and is visualized by GFP::TLN-1 (Walser et al., 2017). During the developmentally regulated L3 turns, TLN-1 is polarized in the direction of the DTC turn (yellow dotted box in Fig. 6C-D, center), where it helps to impart torque that rotates the DTC under pressure from the pushing force of the germline (Agarwal et al., 2022). Before and after turning, we observe that this protein is polarized toward the ventral and dorsal body walls, respectively (blue dotted box, Fig. 6C-D, left and right). We interpret this as indicating the site of interaction that maintains the path of the DTC along these surfaces. The *flh-1/2* double mutant maintains the proper direction of GFP::TLN-1 polarization during the L3 larval stage, and is even slightly more strongly polarized directly after turning (Fig. 6D). Given that *flh-1/2* double mutants do not have turning defects in the L3 stage, it is unsurprising to find that TLN-1::GFP polarization is not disrupted at this stage in the mutants.

### *flh-1/2* prevents cell repolarization in L4

We next examined the localization of GFP::TLN-1 in the DTC of *flh-1/2* mutants (Fig. 7A-B) for aberrant polarization. If the ventral return is associated with the DTC polarizing away from the body wall as it does during the normal L3 turn, it could suggest heterochronic reactivation of the DTC turning program. Indeed, the L4 dorsal DTC enrichment of GFP::TLN-1 is absent in *flh-1/2* double mutants, resulting in roughly equal dorsal and ventral levels (Fig. 7B, C). Since aberrant turns in the L4 occur in under a third of mutants, we further divided our sample into mutants with and without extra turns. In mutants with extra turns, the polarity of GFP::TLN-1 reverses to a ventral enrichment (Fig. 7B, C), while in those with normal migration, GFP::TLN-1 levels are equalized on the dorsal and ventral sides (Fig. 7B, C).

**Figure 7.**
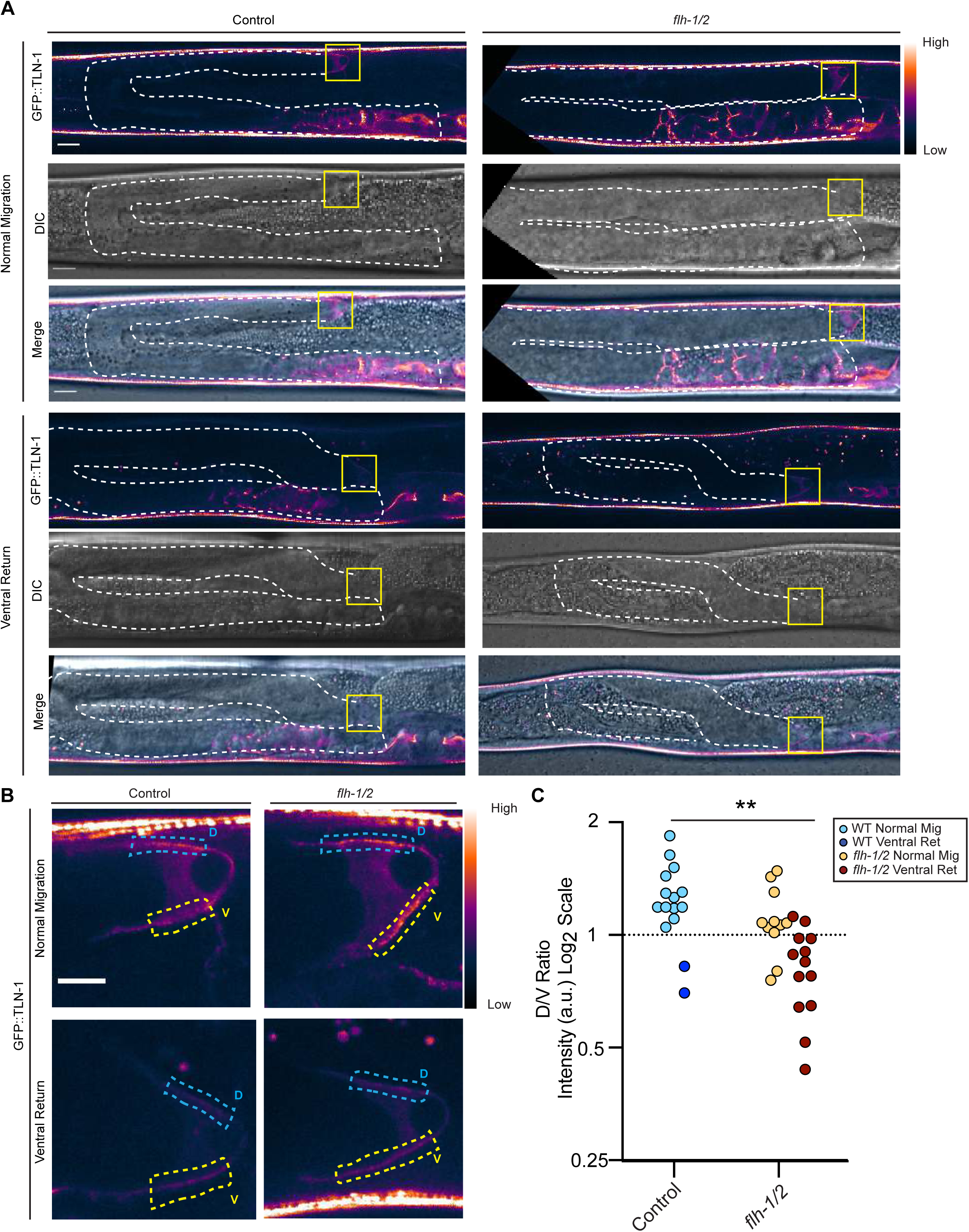
During the ventral return of *flh-1*/2 mutants, GFP::TLN-1 is mispolarized in the direction of turning. (A) Representative images of GFP::TLN-1 animals in otherwise wild-type and *flh-1/2* double mutant backgrounds with examples of both normal migration and ventral return phenotypes with GFP::TLN-1 (top, “Gem” lookup table), DIC (middle), and merged images (bottom). Gonad arms are outlined with white dashed line. DTC regions shown in (B) are marked with a yellow box. Scale bar: 10 μm. (B) Inset images of GFP::TLN-1 localization in L4 DTCs in otherwise wild-type and *flh-1/2* double mutant animals with normal migration (top row) and with the ventral turn phenotype (bottom row). Yellow “A” and blue “B” boxes show ventral and dorsal regions, respectively, of the DTC quantified in (C). Scale bar: 5 μm. (C) Quantification of the ratio of dorsal to ventral intensity (B/A) of GFP::TLN-1comparing *flh-1/2* double mutant animals to wild type. Light blue is otherwise wild-type with normal migration, dark blue is otherwise wild-type with the ventral return phenotype. Yellow is *flh-1/2* with normal migration, dark red is *flh-1/2* with the ventral return phenotype. The dotted line y=1 is given for reference. For wild type n=15, for *flh-1/2* double mutants n=24. Two-tailed unpaired t-test comparing wild types and *flh-1/2* double mutants (**p = 0.0018, t=3.357, df=37).

Remarkably, in two otherwise wild-type animals expressing GFP::TLN-1, we observed aberrant L4 migration in which the DTC made a ventral return; these animals also displayed ventral enrichment of GFP::TLN-1 (Fig. 7B, C, dark blue). The presence of ventrally enriched GFP::TLN-1 in samples of both genotypes that make a ventral return suggests that the extracellular matrix-cytoskeletal linkage via TLN-1 is the mechanism responsible for the extra turns. For DTCs in the *flh-1/2* mutants that have roughly equal dorsal and ventral GFP::TLN-1, the direction of migration appears to be randomized. Because *lin-42* RNAi knockdown in the *flh-1/2* mutants does not rescue the ventral return, there must be other targets of FLH-1/2 in the DTC that ensure proper DTC pathfinding at this stage; we propose some of these will regulate the polarization of TLN-1 in the DTC towards the dorsal body wall.

## DISCUSSION

Our study reveals a role for the FLYWCH transcription factors *flh-1* and *flh-2* as well as *lin-42* in regulating DTC migration during the L4 stage. Through genetic and molecular analyses, we demonstrate that *flh-1*, *flh-2*, and *lin-42* itself protect against L4 DTC turning by dampening the L4 expression peak of *lin-42* and coregulating other targets, thus ensuring the proper localization of TLN-1 toward the dorsal body wall in the final phase of DTC migration. These findings reveal novel functions of *flh-1/2,* acting in a *lin-42*-autoregulatory loop, and integrating the transcriptional control of still-unknown targets with TLN-1 localization to maintain directed cell migration in L4 DTCs.

LIN-42 is not thought to bind DNA directly, but instead via transcriptional co-repressors (Kinney et al., 2023). Indeed, the *lin-42* ortholog *period*/PER is famously autorepressive in *Drosophila* (Hardin et al., 1990) as is Period in mammals (Shearman et al., 2000). If FLH-1/2 act as transcriptional co-repressors with LIN-42–and if this repressive FLH-1/2-LIN-42 complex autorepresses *lin-42* transcription as one of its targets–then we expect to see the results summarized in (Fig. 8). Transcription of *lin-42* will be derepressed in *flh-1/2* double mutants (Fig. 4C), and genetic loss of function of *flh-1/2, lin-42* RNAi, and the *flh-1/2* mutant on *lin-42* RNAi would have the same phenotype (Fig. 5A). The same genetic loss-of-function result could also be obtained if LIN-42 and FLH-1/2 negatively regulate a set of common targets (including the *lin-42* promoter itself) even without directly interacting, if repression of the targets requires the activity of all three repressors to prevent aberrant turning. Since *lin-42* knockdown in *flh-1/2* mutants does not rescue the ventral return phenotype (Fig. 5), we conclude that DTC migration is sensitive to the level of *lin-42* and not simply to the presence or absence of its gene product. We discovered that LIN-42 transcription factor co-regulators known to act in the larval skin, NHR-23 and NHR-85, also bind the *lin-42a* promoter (Fig. 3B), suggesting the capacity for cell-type-specific LIN-42 autorepression.

**Figure 8.**
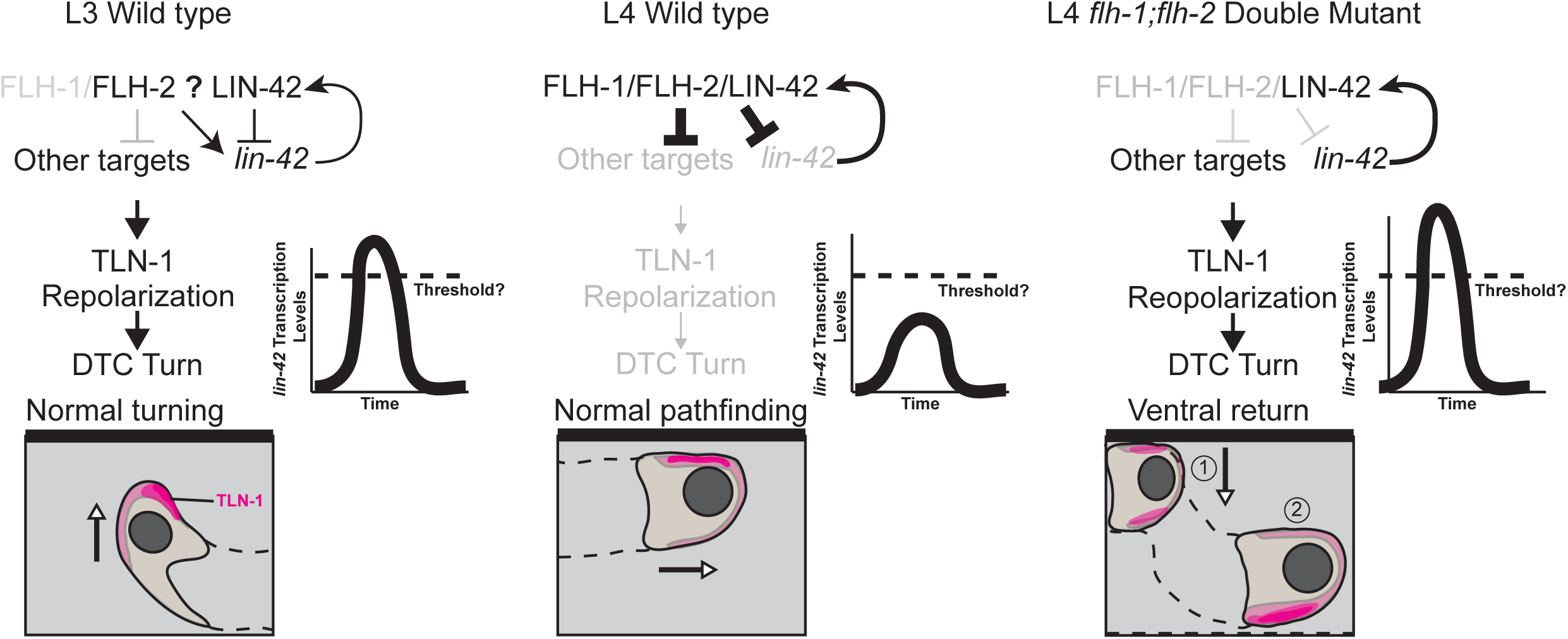
FLH-1/2 and LIN-42 itself repress peak *lin-42* expression and prevent L4 ventral return. The turning phase of distal tip cell migration occurs in a wild type L3 animal (left) in which *flh-1* is not highly expressed, allowing *lin-42* to escape FLH-1/2-mediated autorepression. LIN-42 expression passes a critical threshold and TLN-1 (magenta) polarizes away from the body wall, allowing the DTC to turn. In a wild-type L4 animal (center), the rising transcriptional profile of *flh-1* allows cooperative FLH-1/2-mediated *lin-42* autorepression to dampen the peak expression of *lin-42* below the critical threshold; TLN-1 does not repolarize and the DTC does not turn. In the *flh-1/2* double mutant L4 DTC (right), *lin-42* escapes FLH-1/2-mediated autorepression, its expression rises past the critical threshold, TLN-1 repolarizes off the body wall, and the DTC makes a ventral turn. Other targets may include known targets of LIN-42 in other cell types and stages (for example, many microRNAs), known factors that regulate DTC turning (Wnt and Netrin signaling components, factors regulating cytoskeletal rearrangement and interaction with the extracellular matrix), and new members of this regulatory pathway yet to be discovered.

Our work contributes to the growing understanding of transcriptional control of DTC migration, which is known to involve transcription factors HLH-2, HLH-12 (Sallee et al., 2017; Meighan et al., 2015; Meighan and Schwarzbauer, 2007; Tamai and Nishiwaki, 2007), LIN-32 (Littleford et al., 2021), and VAB-3 (Meighan and Schwarzbauer, 2007). How a repressive complex of FLH-1/2-LIN-42 might interact with these factors, or whether LIN-42 can inhibit their binding to cognate promoters, should be investigated.

The biology of the *lin-42* locus is complex (Edelman et al., 2016; Monsalve et al., 2011; Tennessean et al., 2006). It is known that rare surviving escapers carrying a *lin-42(ox460)* or *lin-42(ox461)* null allele can sometimes, instead of arresting at an early larval stage, form a gonad that begins to grow and even undergo turning (Edelman et al., 2016). These gonads are extremely small, with very few germ cells. Germline size is reduced in mutants carrying a different allele, *lin-42(n1089)*, likely because *lin-42* promotes the expression of the stem cell niche cue, *lag-2,* in the DTC (Berardi et al., 2018). Investigation into how the various functions of *lin-42*, including in DTC migration, are spread across its complex regulatory and protein coding regions is ongoing. We also aim to integrate our findings with other timing factors, such as LIN-29 and BLMP-1, known to operate in the DTC turning regulatory network (Ambros and Horvitz, 1984; Huang et al., 2014; Stec et al., 2021; Tennessen et al., 2006).

Further work is needed to explore genetic underpinnings of the L4 ventral return vs. the normal L3 turn. We hypothesize that the ventral return may be a heterochronic reactivation of the L3 turning program. In both the L3 turn and the ventral return, *lin-42* is at its peak of expression and TLN-1 polarizes off the body wall along which the DTC had previously been migrating. FLH-1/2 repression in the wild-type L4 would therefore act to repress the entire turning program at this later stage. What additional interactions explain the switch between L3 activating and L4 repressing regulation of *lin-42* by FLYWCH factors? In the whole-body transcriptome study, *flh-1* expression rises ∼5-fold from the L3 baseline through early L4 (Fig. 3 “rising”, Fig. S4C reanalyzed data from (Hendriks et al., 2014)). We hypothesize *lin-42* is not repressed during the L3 turn by FLH-1/2 because *flh-1* expression is too low at that stage, and FLH-2 alone does not bind the *lin-42a* promoter (at least not in our Y1H). In this model, FLH-2 would act, either indirectly or with a different binding partner, as an activator of *lin-42* in L3 that switches to a repressor in L4 as FLH-1 accumulates in the DTC. We also must align our finding that the L3 turn occurs in a “*lin-42* ON” transcriptional state with the fact that *lin-42* RNAi leads to precocious turning in L2 (Fig. S2) (Tennessen et al., 2006). We find genetic and transcriptional evidence of autoregulatory repression of *lin-42* transcription, which complicates the inference of a factor being “on” or “off” in a given cell at a given time based on transcriptional activity of its promoter–it may well be that transcriptional “on” state for *lin-42* corresponds to low levels of LIN-42 protein and vice versa, as is the case for the orthologous *Drosophila* gene *period* (Hardin et al., 1990). This may explain the difference between LIN-42::YFP dynamics and *lin-42a* promoter dynamics. One caveat is that the *lin-42a* and *lin-42b* promoter fusions also lack the *lin-42* endogenous 3’ UTR, meaning they are not subject to the same post-transcriptional regulation as *lin-42* itself.

MicroRNAs could mediate such post-transcriptional regulation. LIN-42 is a key regulator of microRNAs broadly (McCulloch and Rougvie, 2014; Perales et al., 2014; Van Wynsberghe et al., 2014), and the only previously known role of FLYWCH factors in *C. elegans* is inhibiting embryonic expression of microRNAs like *lin-4* (Ow et al., 2008). Although *lin-4* mutants have normal DTC migration (Fig. 1G), *lin-4* should be investigated for potential redundancy with its MIR-10 family members *mir-51, mir-57,* and *mir-237* (Clarke et al., 2025; Fromm et al., 2015). Both *lin-4* and *mir-237* have an identical seed sequence (Clarke et al., 2025) and rank among the most overexpressed microRNAs in *lin-42(n1089)* loss-of-function mutants at the L4 stage (Van Wynsberghe et al., 2014). The most strongly derepressed microRNA at the L4 stage in this dataset, *let-7,* is a known target of *lin-42* at earlier larval stages (McCulloch and Rougvie, 2014). Whether FLH-1/2 act alongside LIN-42 to suppress its other target microRNAs in embryos or later stages remains an open question. Finally, known regulators of DTC turning robustness, *mir-34* and *mir-83* (Burke et al., 2015), are also strong candidates for LIN-42 repression of L4 turning. They are both overexpressed (∼1.2x and ∼1.5x, respectively) in *lin-42(n1089)* mutants at the L4 stage (Van Wynsberghe et al., 2014). *lin-42(n1089)* affects only the *lin-42b* and *lin-42c* isoforms, so it is possible that *lin-42a* might repress a distinct subset of microRNAs. Notably, the pattern that we identified here–a transcription factor that represses peak expression levels of a target and is expressed in phase with that target (*flh-2* with *lin-42*) is similar to the pattern observed for *lin-42* and its target *let-7* microRNAs in L1 and L2 larvae (McCulloch and Rougvie, 2014).

MicroRNA-mediated mRNA instability plays a particularly important role in generating oscillations in gene regulatory networks, especially when combined with autorepression (Minchington et al., 2020). More generally, we consider that proper developmental control involves not only the presence or absence of regulatory factors, but also their appropriate relative dosage. Autorepression can also dampen noise in gene regulatory circuits (Okano et al., 2008). During the L4 stage, *lin-42* is expressed at relatively low levels in the DTC compared to other cell types (Fig. 1, 2). However, this means that its derepression in *flh-1/2* double mutants causes a dramatic change in its relative expression levels (Fig. 4). One might expect that genes expressed at low levels have particularly strong dosage-dependent effects. For example, in *Drosophila* blastoderm embryos, bimodal readouts of gene regulatory gradients are highly sensitive at the edge of a concentration threshold (Crocker and Erives, 2013).

There is considerable diversity in gonad morphology among nematode species. *Pristionchus pacificus* hermaphrodite gonads are didelphic, yet both gonad arms display a ventral return during the late fourth larval stage (J4) in both gonad arms resulting in a “pretzel-shaped” morphology (Rudel et al., 2005; Pires-daSilva & Parihar, 2012). It would be interesting to explore whether this regulated ventral return shares molecular similarity with the *C. elegans* ventral return mutant phenotype we characterized here.

Notably, while *Pristionchus pacificus* has a *lin-42* homolog, it lacks clear FLH-1 and FLH-2 homologs; their best BLASTP match is a single gene, sequence PPA10311 with which they share 21% and 18% sequence similarity, respectively. In the developing *C. elegans* male gonad the linker cell guides the growth of the single gonad arm, which makes an early turn from ventral to dorsal, followed by a ventral return (though these turns occur at the L2 and L3 stages, respectively) (Kimble & Hirsh, 1979). *flh-1* and *flh-2* are expressed in the male linker cell, where there is no evidence of *lin-42* expression (data from Schwarz et al., 2012). Future work should investigate whether the gene regulatory pathway identified in the DTC also contributes to regulating the timing of migration of the male linker cell and DTCs of other nematode species.

The FLYWCH zinc-finger-containing transcription factors are themselves noteworthy. FLYWCH transcription factor genes are *Phantom* elements derived from the *Mutator* DNA transposase and are present in many eukaryotic genomes (Babu et al., 2006; Marquez and Pritham, 2010). Although FLYWCH transcription factor sequences are homologous to one another, they have independently expanded in metazoan transcription factor repertoires several times (Mukherjee and Moroz, 2023).

*C. elegans* FLYWCH transcription factors are homologous to the FLYWCH transcription factors of *Drosophila melanogaster* and human FLYWCH1/KIAA1552 (Babu et al., 2006). In human colorectal cancer, FLYWCH1 acts as a novel repressor of Wnt/β-catenin signaling, by which it regulates cell polarity and migration (Muhammad et al., 2018). Wnt signaling is the primary regulator of *C. elegans* DTC polarization and migration along the anterior/posterior axis (Levy-Strumpf et al., 2015; Levy-Strumpf and Culotti, 2014), suggesting either a convergent or deeply conserved regulation of the Wnt-responsive cell migration polarity by animal FLYWCH transcription factors. Future research will explore the relationship between *flh-1/2* and Wnt-mediated cell polarity in the DTC. The role of these FLYWCH factors in *lin-42* regulation is remarkable for its cell-type and temporal specificity, despite the fact that *lin-42* transcription is synchronized across many cell types (Monsalve et al., 2011; Hendriks et al., 2014; Tennessen et al., 2006). This suggests a paradigm by which global synchronization of gene expression activation by a broadly expressed regulator (yet to be discovered) can be modulated by cell-type-specific transcription factors to allow the progression of stage-specific development.

## MATERIALS AND METHODS

Sections of this text are adapted from (Li et al., 2022; Singh et al., 2024), as they describe our standard laboratory practices and equipment. Reagents are listed in Table S1. Reagent Table.

### C. elegans strains

*C. elegans* strains were maintained on NGM plates seeded with OP50 *E. coli* at 20°C. Some strains were provided by the *Caenorhabditis* Genetics Center (CGC), which is supported by the National Institutes of Health - Office of Research Infrastructure Programs (P40 OD010440). In strain descriptions, we designate linkage to a promoter with a p following the gene name and designate promoter fusions and in-frame fusions with a double colon (::).

### Yeast strains

Yeast strains were maintained using YPD as previously described (Deplancke et al., 2006; Reece-Hoyes et al., 2011).

### Yeast one-hybrid assays

Yeast one-hybrid assays were performed by Gibson Assembly of a bait target site into the pMW2 yeast one-hybrid vector, transformed into the YM4271 yeast one-hybrid strain, which was subsequently mated to the wTF2.2 gal4 AD transcription factor library (Deplancke et al., 2006). Protein-DNA interactions were scored by visible growth on SC-HIS -TRP plates with 3-AT compared to empty vector controls after three days of growth by both a 1:200 dilution in ddH20 and by streaking with a toothpick. Images were created by plating 3 μL drops of 1:200-diluted overnight cultures and imaged on a Bio-Rad Universal Hood II gel imager. Images were inverted on Image Lab Software (RRID:SCR_014210). Because BLMP-1 has been shown to be both a pioneer factor for heterochronic genes (Stec et al., 2020), and a repressor of DTC turning genes (Huang et al., 2014), target bait sites were selected based on sites of BLMP-1 ChIP seq peaks upstream of the target gene found on ModENCODE (Araya et al., 2014) tracks on the *C. elegans* WormBase JBrowse genome browser (Diesh et al., 2023).

### RNAi Feeding

HT115(DE3) E. coli from the Ahringer (Kamath, 2003) and Vidal unique (Rual et al., 2004) RNAi libraries with or without (empty vector control L4440) the dsRNA trigger insert for the intended target gene was grown overnight at 37°C and induced to express dsRNA by exposure to 1 mM IPTG for one hour at 37°C. RNAi bacteria was plated on NGM plates with IPTG and Ampicillin. Animals were placed on RNAi-seeded plates as L4s and F1 progeny were used for imaging.

For the construction of the *lin-42* RNAi plasmid, exons 6-9 were amplified from pCMH1968 and cloned by Gibson Assembly into vector L4440 using primers 5’-CCAGTACCATCCACCTCCCGTCACA-3’ and 5’-TTAATTCTGAGAATCCCGTAGCATT-3’. The construct was sequenced for correctness and worm exposures were conducted by RNAi feeding as described above. Worms were examined for precocious alae phenotype, precocious DTC turning, and knockdown of LIN-42::YFP to confirm knockdown efficiency (Fig. S2).

The DTC-specific RNAi strain NK211543 (Linden et al., 2017) lacks global RNAi activity due to a *rde-1(ne219)* loss-of-function allele rescued in *lag-2p*-expressing cells by the *lag-2p::mNG::PLC^δPH^::F2A::rde-1* transgene that also labels the DTC, and a *rrf-3(pk1426)* mutation that enhances RNAi activity. Experiments with this strain are conducted at a temperature of 16°C.

Generation of lin-42ap::SV40-NLS::GFP::PEST; lin-42bp::SV40-NLS::mCherry::PEST strain 3,048-bp upstream of the *lin-42a* start site (5’-AGAATTCGGGAAAAGGACCAGAAGG to 5’-CTAAGCCTACCACCCTAAAATGGAT-3’) and 4,210-bp upstream of the *lin-42b/c* start site (agctaattttcaagctcctactaga…cagaacgtggcaccatcagccaaat) were amplified from *C. elegans* N2 genomic DNA. The *lin-42a* fragment was subcloned into pCMH1225 (*SV40-NLS::GFP::pest::unc-54*) upstream of *NLS::GFP::PEST* to make pCMH1207. The *lin-42b/c* fragment was subcloned into pCMH1183 upstream of *NLS::mCherry::PEST* to make pCMH1210. Both plasmids along with plasmids containing *unc-119(ed3)* and *ttx-3*::GFP were co-injected into *unc-119(lf)* animals. The extrachromosomal arrays were integrated by gamma-ray mutagenesis and the strain was backcrossed to N2 five times.

### Confocal microscopy

All images were captured at room temperature using a Leica DMI8 with an xLIGHT V3 confocal spinning disk head (89 North), equipped with a 63× Plan-Apochromat (1.4 NA) objective and an ORCAFusion GenIII sCMOS camera (Hamamatsu Photonics) controlled by microManager (Edelstein et al., 2010). RFPs were excited with a 555 nm laser, and GFPs with a 488 nm laser. Z-stacks through the gonad were acquired with a step size of 1 µm, except for the TLN-1::GFP localization experiments (see below). Worms were mounted on agar pads using 100 mM sodium azide.

### *C. elegans* larval development staging

L3 animals were staged based on the number of P6.p cells (1 = early, 2 = middle, and 4 = late) in Figure 1 and on migration of the DTC (preturn, turning, post-turn) in Figures 2, 6, S5, & S6. L4 animals were staged according to vulval morphology outlined in (Mok et al., 2015).

### Fluorescence intensity quantification of nuclei

Images were processed in ImageJ/FIJI (Schindelin et al., 2012). Cell types were identified with DIC optics. Intensity was quantified by measuring the mean gray value (in arbitrary units) of a region of interest and subtracting from it the mean gray value of a background region inside the worm. For *lin-42ap::GFP*, LIN-42::YFP, and *lin-42bp::mCherry* images were taken at laser power 50% with 200 ms exposure. NHR-85::GFP images were taken at 10% laser power with 500 ms exposure time. NHR-23::mScarlet images were taken at 50% laser power at 500 ms exposure. For representative images in the figures, those that required stitching were stitched using the ImageJ Stitching Plugin (Preibisch et al., 2009) and straightened using the straightening tool.

### Quantifying GFP::TLN-1 localization

To quantify GFP::TLN-1 localization in the DTC for Figures 6 and 7 we selected specimens at the correct stage based on vulval morphology (Mok et al., 2015) and DTC position. Confocal Z-stacks acquired at 0.3 μm z-steps with a laser power of 50% and an exposure time of 200 ms using the 488 nm laser. A central z-slice with both sides of the DTC in cross-section was selected from each specimen. GFP::TLN-1 was measured using a 1 pixel-wide x ∼2-10 micron long line drawn through the cross section of the cell region of interest with the line tool in FIJI/ImageJ (Schindelin et al., 2012).

The line segment was moved to take a background measurement in the center of the cell (a void in cross-section), and this background measurement was subtracted from the GFP::TLN-1 measurement.

### Graphing and statistical analysis

Graphs were generated using GraphPad Prism (GraphPad Prism Version 10.10 (264) for macOS), GraphPad Software, Boston, Massachusetts USA). Sample sizes, tests, test statistics, and p values are given for each analysis in the accompanying figure legends. Statistical testing and corrections for multiple comparisons as described in the figure legends were carried out using GraphPad Prism and R using the statix package (Kassambara, 2023).

### Transcriptome re-analysis

SRA data acquired from Hendriks et al. 2014 (GEO accession number GSE52910) for developmental (contDevA) samples ranging from 21 hours to 36 hours (Hendriks et al., 2014). Downloaded RNA-seq reads were aligned to the latest available *C. elegans* genome from WormBase (WBcel235)(Harris et al., 2019). Alignments and a count table were generated using the R package QuasR (www.bioconductor.org/packages/2.12/bioc/html/QuasR.html). Initial read alignment was performed using the built-in Hisat2 (http://daehwankimlab.github.io/hisat2/) alignment algorithm. The primary command used to generate alignments was “align<-qAlign(sampleFile = “samples.txt”,genome = “ce.fa”,splicedAlignment = TRUE,aligner = “Rhisat2”). Individual exon counts were then obtained by using the quantify alignment command “(counts<-qCount(proj = align,query=exons,orientation = “opposite”,selectReadPosition = “end”,reportLevel = “exon”). We accounted for variability in library depth using a similar normalization method as the authors of Hendriks et al., 2014. Each sample was divided by the total number of reads and then multiplied by the total average library size to normalize library size across all samples. After normalization, the count table was used to measure expression levels of both the *lin42a* and *lin42b* isoforms. To determine the expression levels of each specific isoform, we measured the expression of exons specific to *lin42a* and *lin42b* independently and compared these expression values. Isoform-specific exons were identified by matching up exon start and end ranges with the corresponding exonID numbers from the WBcel235 annotation file (gff3). For this analysis, *lin42a* was assigned exonID number 39449, while the *lin42b* isoform was tracked by expression of exonID numbers 39443, 39446, 39447, 39448, 39453, 39454, and 39455. Shared exons were assigned exonID numbers 39443, 39446, 39447, and 39448. The *flh-1* gene has three annotated isoforms, two of which have a small (17nt) first exon transcribed from an alternative transcription start site, so we focused on the shared exons which make up the shortest isoform, *flh-1c*. These exons were assigned exonID numbers 95486, 95487, and 95488.

## ACKNOWLEDGEMENTS

The authors would like to thank Dr. Victor Ambros for sharing strains. We thank Camille Miller for technical support and Dr. Taylor Medwig-Kinney, Noor Singh, Xin Li, Bailey de Jesus, and five reviewers for helpful feedback. We thank Dr. Rob Dowen and Peter Breen for sharing RNAi clones and Dr. Kerry Bloom for use of his lab space and sharing yeast reagents. We would also like to thank the *Caenorhabditis* Genetics Center which is funded by NIH Office of Research Infrastructure Programs (P40 OD010440).

## DATA AND RESOURCE ACCESSIBILITY

The authors affirm that the data necessary for confirming the conclusions of the article, with the exception of Figure S4, are present within the article, figures, and tables. Strains and plasmids are available upon request. The data necessary for confirming Figure S4 is from Hendriks et al., 2014, Accession Number: GSE52910.

## SUPPLEMENTAL INFORMATION

**Figure S1.** Loss of FLYWCH gene function causes DTC migration defects. (A) Representative image of normal gonad migration. Scale bar: 10 μm. (B) Representative image of an animal categorized as “ventral return.” (C) *flh-2(bc375)* animal categorized as “dorsal turn”. Gonad migration is initially ventralized with a later dorsal turn (n= 2/32). The gonad is outlined with a dashed white line. The DTC is marked with a yellow asterisk. (C) Percentage of samples with each phenotype for genotypes shown at bottom. Wild type, *flh-1(tm2118),* and *flh-2(tm2126)* and the double mutant *flh-1(tm2118); flh-2(tm2126)* are the same datasets as in Fig. 4B. (Wild type n=50, *nhr-85(ok2051)* n=33, *nhr-23* RNAi n=30, *flh-1(bc374)* n=32, *flh-2(bc375)* n=32, *flh-1(tm2118)* n=31, *flh-2(tm2126)* n=32, *flh-3(gk1049)* n=32, *flh-1/2* double mutant n=52, *flh-1/2* double mutant on *flh-3* RNAi n=32). The *flh-3(gk1049)* allele (a 1,200-bp deletion of the first two exons) resulted in a 18.75% defect rate while *flh-3* RNAi-treated wild type animals had a 13.33% ventral return defect rate. Statistical analysis was performed using a pairwise proportion test, with p-values adjusted for multiple comparisons via the Benjamini-Hochberg procedure, which showed that differences were not significant.

**Figure S2.** *lin-42* RNAi achieves efficient knockdown. (A) Skin and developing vulva of an early L4 animal fed control and *lin-42* RNAi. The white arrow points to precocious alae (n=4/10 for *lin-42* RNAi treated animals and n=0/10 for controls), cuticular features that appear at the wrong time after *lin-42* loss of function (Banerjee et al., 2005; Tennessen et al., 2006). Scale bar: 5 μm. (B) Early DTC turns observed after whole-body *lin-42* RNAi (n=3/30) and control RNAi fed animals (n=0/30) at early L3. Stage determined by vulval precursor cell number (white arrow). The gonad arm is outlined with a white dashed line. Scale bar: 5 μm (C) Early DTC turns observed after DTC-specific *lin-42* RNAi (n=2/31) and control RNAi fed animals (n=0/30) at early L3. Stage determined by vulval precursor (white arrow) cell count. The gonad arm is outlined with a white dashed line. Scale bar: 5 μm (D) Control and *lin-42* RNAi fed LIN-42::YFP stage L4.4-4.5 animals at the peak of LIN-42 expression with (right) and without (left) DIC merge showing hyp7 cells (arrows) quantified for (E). Scale bar: 5 μm. (E) Quantification of LIN-42::YFP hyp7 fluorescence intensity comparing control (n=10) and *lin-42* RNAi (n=13) in L4.4-4.5 hyp7 cells. Each data point represents an average of three hyp7 cell measurements in one animal. Mixed-effects model (REML) F _1, 21_ = 24.33 (****p <0.0001).

**Figure S3.** Transcriptional reporters of two *lin-42* promoters are dynamically expressed across cell types in L4. (A) Graphical representation of the *lin-42* locus and design of *lin-42ap::SV40-NLS::GFP::PEST*; *lin-42bp::SV40-NLS::mCherry::PEST* transcriptional reporter, each fused to a fluorescent protein coding gene with an SV40-NLS. Because of the dynamic expression we observed for LIN-42::YFP, we included PEST tags on both fluorescent proteins to promote turnover and capture signal from only newly synthesized proteins (B) Schematic illustrating the locations of the different cell types analyzed. (C) Representative images taken at L4.5 for *lin-42ap::SV40-NLS::GFP::PEST* and L4.7 for *lin-42bp::SV40-NLS::mCherry::PEST*. Both *lin-42a* and *lin-42b* are expressed in the DTC (purple), seam cells (blue), hyp7 cells (orange), and vulval cells (red); colored dotted boxes designate focal cells in each panel. Scale bar: 5 μm. (D) Quantification of intensity of *lin-42ap::SV40-NLS::GFP::PEST* and *lin-42bp::SV40-NLS::mCherry::PEST* expression in seam cells, hyp7 cells, vulval cells, and DTC in each L4 substage. Each data point represents the average fluorescence intensity from individual animals measured by the mean intensity of three cells for each cell type except for DTC which was measured one per animal (n=8-12 per substage). Dotted lines represent SEM. In most tissue types, the temporal discrepancy between the two peaks of transcriptional activity is greater than what could reasonably be explained by differences in GFP and mCherry folding times (Balleza et al., 2018).

**Figure S4.** *lin-42* transcripts in populations of worms oscillate synchronously in the L3-L4 stages, while *flh-1c* rises. (A) A graph depicting average read number and SEM comparing a *lin-42a* specific exon (green), *lin-42b* specific exon (magenta), and exon shared between the two isoforms (gray) from bulk RNA-seq Illumina reads (Hendriks et al., 2014) from synchronized animals reared at 25°C, collected at one-hour intervals starting with 21 hours post-release from L1 synchronization until 36 hours post-release from L1 synchronization. (B) Data from (A) comparing *lin-42a* specific exon (green) and *lin-42b* specific exon (magenta) only. All error bars denote SEM. (C) A graph depicting average read number and SEM of *flh-1c* exons from Illumina sequencing results (Hendriks et al., 2014) from synchronized animals at one hour intervals starting with 21 hours post-release from L1 synchronization until 36 hours post-release from L1 synchronization. This bulk RNA-seq data will not reflect any DTC-specific expression dynamics, since the DTCs are a small minority of cells and these genes are not specifically expressed in the DTC.

**Figure S5.** *lin-42* is autorepressive in the larval skin in L4. (A) Expression intensity of *lin-42ap::SV40-NLS::GFP::PEST* and (B) *lin-42bp::SV40-NLS::mCherry::PEST* in control and lin-42 RNAi treated mutant animals in the DTC during preturn L3, turning L3, post turn L3 and each L4 substage. Each data point represents the average fluorescence intensity from individual animals measured by average of three cells for each cell type except for DTC which was measured one per animal (n=7-12 per substage). Dotted lines represent SEM. Two-way ANOVA with Tukey’s multiple comparisons test for *lin-42a* (F (27, 217) = 14.91) and *lin-42b* (F (27, 217) = 6.010) (****p <0.0001) (***p <0.001). (C) Expression intensity of *lin-42ap::SV40-NLS::GFP::PEST* in L4.5 hyp7 cells comparing wild-type animals on control RNAi (gray, n=10), wild-type animals on *lin-42* RNAi (blue, n=16), *flh-1/2* double mutants on control RNAi (red, n=18), and flh-1/2 double mutants on *lin-42* RNAi (yellow, n=16). Two-way ANOVA with Tukey’s multiple comparisons test (F (3, 33) = 15.36)(****p <0.0001)(***p <0.001)(**p <0.01) (*p <0.05).

**Figure S6.** The *lin-42b* promoter is also regulated by *flh-1/2*. (A) Expression intensity of *lin-42bp::SV40-NLS::mCherry::PEST* in wild type and *flh-1/2* double mutant animals in the DTC, seam cells, hyp7 cells, and vulval cells in each L4 substage. Each data point represents the fluorescence intensity from an individual animal measured as the mean intensity of three cells per cell type (seam, hyp7, vul), or intensity of a single DTC per animal (n=8-12 per substage). Statistics were performed using a Two-way ANOVA with Tukey’s multiple comparisons test on the DTC (F (21, 183) = 15.45), seam cells (F (21, 184) = 20.55), hyp7 cells (F (21, 184) = 20.74), and vulva (F (21, 184) = 15.76)(*p <0.05)(**p <0.01)(***p <0.001)(****p <0.0001). (B) Representative images from three staged specimens each of *lin-42bp::SV40-NLS::mCherry::PEST* in otherwise wild-type controls. The gonad is outlined with a white dotted line, and the DTC is marked with a white arrow. Scale bar: 5 μm. (C) Quantification of fluorescence intensity of *lin-42bp::SV40-NLS::mCherry::PEST* in otherwise wildtype (gray), and *flh-1/2* (purple) during each L3 DTC turning stage. Values are normalized to post-turn mean expression for each genotype. Two-way ANOVA with Tukey’s multiple comparisons test (F (5, 37) = 7.442) (*p <0.1). (D) Expression intensity of *lin-42bp::SV40-NLS::mCherry::PEST* in control and *lin-42* RNAi treated animals in the DTC during preturn L3, turning L3, post turn L3 and each L4 substage. Each data point represents the average fluorescence intensity from individual animals measured by average of three cells for each cell type except for DTC which was measured one per animal (n=7-12 per substage). Dotted lines represent SEM. Statistics were performed using a Two-way ANOVA with Tukey’s multiple comparisons test (F (27, 203) = 2.676), and found no significant difference of *lin-42b* expression in the DTC after *lin-42* RNAi.

**Figure S7.** Yeast one-hybrid hits from the *lin-42b* promoter screen. (A) A graphical representation of the *lin-42* locus and the region used as bait used for the *lin-42b* yeast one-hybrid screen. (B) List of *lin-42b* bait-binding transcription factor genes, including their dynamic transcription factor class and phase (from Hendriks et al., 2014).

---

**Table S1.** Reagent Table.

